# The future of a partially effective HIV vaccine: assessing limitations at the population level

**DOI:** 10.1101/509828

**Authors:** Christian Selinger, Dobromir T. Dimitrov, Philip Welkhoff, Anna Bershteyn

## Abstract

**Objectives:** Mathematical models have unanimously predicted that a first-generation HIV vaccine would be useful and cost-effective to roll out, but that its overall impact would be insufficient to reverse the epidemic. Here, we explore what factors contribute most to limiting the impact of such a vaccine.

**Methods:** Ranging from a theoretical ideal to a more realistic regimen, mirroring the one used in the currently ongoing trial in South Africa (HVTN 702), we model a nested hierarchy of vaccine attributes such as speed of roll-out, efficacy, and retention of booster doses.

**Results:** The predominant reasons leading to a substantial loss of vaccine impact on the HIV epidemic are the time required to scale up mass vaccination, limited durability and waning of efficacy.

**Conclusions:** A partially effective HIV vaccine will be a critical milestone for the development of a highly effective, durable, and scalable next-generation vaccine. Accelerated development, expedited vaccine availability, and improved immunogenicity are the main attributes of a vaccine that could dramatically reverse the course of the epidemic in highly endemic countries.

## 1 Introduction

An estimated 2.1 million people were infected with HIV in 2015 [3]. Despite increasing numbers of people on antiretroviral treatment (ART), there is still a need to scale up HIV prevention in order to counter the global epidemic on a population level. Existing prevention modalities such as condoms, medical male circumcision, treatment as prevention, and oral pre-exposure prophylaxis (PrEP) face limitations such as negotiability, stigma, access, adherence, retention, and efficacy [30, 10]. A breakthrough in HIV prevention such as a highly effective vaccine is urgently needed. The Pox-Protein Public-Private Partnership (P5) is working to build on the findings of the RV144 trial [34] with a currently ongoing Phase 2b/3 trial (HVTN 702) in South Africa [1]. With its complex immunization schedule and anticipated waning of immunity, the regimen with currently 6 doses potentially followed by frequent boosts is likely to provide only partial efficacy over limited time. Such vaccine could be seen as a first-generation product that must still be improved upon in order to fundamentally transform the HIV epidemic. Mathematical models [23, 19, 39, 6, 18, 26, 25, 16, 31, 5, 24] suggest that the first-generation vaccine would be useful and cost-effective to roll out, but that its overall impact will be modest. Here, we explore what factors contribute collectively to limiting the impact of such a vaccine.

## 2 Methods

We developed an agent-based model of the South Africa population to forecast HIV infections over a 20-year time horizon, from year 2027 to 2047. As compared to a reference case with no HIV vaccine, we evaluate implementation of strategies for initiation of HIV vaccination.

### Model set-up and calibration

We modified EMOD-HIV v2.5, an age-stratified and individual-based network model of HIV of South Africa, to incorporate HIV vaccination according to pox-protein HIV vaccine regimens (such as the regimen currently being tested in HVTN 702). Because EMOD is an individual-based model, interventions such as a time-varying course of vaccine efficacy can be applied to each individual according to his or her own timing of vaccination and adherence to the booster series. This renders the model well suited for a nuanced analysis of the anticipated time-dependent efficacy of the pox-protein HIV vaccine regimen.

The parameters, model input values, sources, projections, and sensitivities of the epidemic projection without vaccine, used as the reference group for comparison, have been described previously [9, 21, 8]. A detailed model description, user tutorials, model installer, and source code are available for download at http://idmod.org/software. EMOD-HIV is an individual-based model that simulates transmission of HIV using an explicitly defined network of heterosexual relationships that are formed and dissolved according to age- and risk-dependent preference patterns [20]. The synthetic population was initiated in 1960, and population recruitment and mortality was assumed to be proportional following age- and gender-stratified fertility and mortality tables and projections from the 2012 UN World Population Prospects [4]. Since the population size of South Africa exceeds the computational limit of simulated agents, we assumed that one simulated agent corresponds to 300 real-world individuals. The model was calibrated to match retrospective estimates of age-stratified, national-level prevalence, incidence, and ART coverage from four nationally representative HIV surveys in South Africa [38, 14, 32, 37, 36]. For each simulated vaccine scenario, we used the 50 most likely parameter sets obtained from the gradient-descent based calibration process. The age patterns of sexual mixing were configured to match those observed in the rural, HIV-hyperendemic province of KwaZulu-Natal, South Africa [29]. Recently, a validation study showed that self-reported partner ages in this setting are relatively accurate, with 72% of self-reported estimates falling within two years of the partners actual date of birth [17]. Further, the transmission patterns observed in EMOD [9] are consistent with those revealed in a recent phylogenetic analysis of the age/gender patterns of HIV transmission in this setting [28].

Transmission rates within relationships depend on HIV disease stage, male circumcision, condom usage, co-infections. Viral suppression achieved through antiretroviral therapy [7, 22] is assumed to reduce transmission by 92%-an estimate based on observational data of serodiscordant couples in which outside partnerships could have contributed to HIV acquisition [12].

### HIV Treatment and Prevention

We configured the EMOD health care system module to follow trends in antiretroviral therapy (ART) expansion in South Africa. Treatment begins with voluntary counseling and testing (VCT), antenatal and infant testing, symptom-driven testing, and low level of couples testing. The model includes loss to follow-up between diagnosis and staging, between staging and linkage to ART or pre-ART care, and during ART or pre-ART care [27]. Projections of South Africa treatment expansion in the no vaccine reference group are calibrated to reflect a gradual decline of HIV incidence without elimination, so that HIV remains endemic through 2050 [13]. All scenarios included medical male circumcision [11] at 22% coverage and conferring 60% reduction in acquisition risk with lifetime durability. Condom usage was dependent on four relationship types (transitory, informal, marital, commercial), with per act usage probability ramping up to median values of 62%, 39%, 26%, and 85% by 2027 across parameter draws. To allow for maximum hypothetical impact of a vaccine, we assume future ART coverage of about 60% (consistent with the current guidelines assumed in the HIV/TB Investment Case Report for South Africa [2]) and no use of oral PrEP. Variations on these assumptions, explored elsewhere [35], only further diminish the impact of the vaccine.

### HIV Vaccine Efficacy

We incorporated a parametric model of time-dependent vaccine efficacy that was hypothesized for the pox-protein regimen based on results from RV144. We included the time series of efficacy associated with each of the originally planned 5 doses administered during the study and possible booster doses (described in detail below) beyond the 24-month duration of the first stage of the study. Specifically, the pox-protein dosing schedule that was modeled consisted of a series of five immunizations over 12 months ALVAC-HIV-C was administered at months 0 and 1, followed by ALVAC-HIV-C+gp120 at months 3, 6, and 12 (supplementary to RV144 schedule). The protocol was later revised to include a boost at month 18, but this was not considered at the time of this modeling effort. Time-dependent vaccine efficacy was interpreted as a per exposure reduction in the probability of acquisition parameterized by an impulse and exponential decay model.

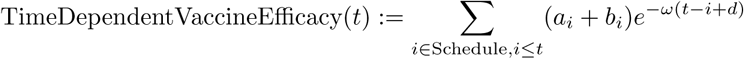

where *a*_*i*_ is the efficacy increase of immunization with ALVAC-HIV-C, *b*_*i*_ is efficacy increase after ALVAC-HIV-C + gp120 immunization, *ω* is the efficacy decay rate per month and *d* is the delay between immunization and initiation of protective effect in months.

Assuming uniformly distributed exposure over a given time span in the trial, we calculated the cumulative vaccine efficacy (corresponding to the efficacy estimate from the trial) as the area under the curve of the instantaneous vaccine efficacy rescaled by the length of the time span.

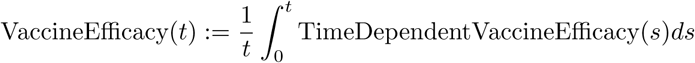

In anticipation of efficacy results for HVTN 702, we modeled time-dependent vaccine efficacy based on results from statistical models [33] for RV144 study outcomes using a point-estimate of 58% shortly after the month 6 vaccination, and cumulative efficacy of 31.2% over 42 months. We adjusted the parameters of the efficacy function such that the cumulative vaccine efficacy over 24 months after the first dose is 50% and obtained values *a*_*i*_ = 0.08, *b*_*i*_ = 0.34, *ω* = 0.065 and *d* = 0.1.

### Booster Schedule and Efficacy

For the purpose of model projections beyond the primary trial endpoint, we also implemented up to four two-yearly boosters starting at month 36 with fixed attrition rates of 0, 20 or 50% per booster to cover a total of 10 years of vaccine efficacy. We assumed booster efficacy to follow the same parameterization as ALVAC-HIV-C + gp120 doses from the primary immunization series during the first 12 months. Booster eligibility depended on having received the primary immunization series or the booster previously. Missing a booster resulted in loss of eligibility for subsequent boosters, which may or may not prove to be the case when the vaccine is implemented. Individuals who tested HIV positive were not eligible for future boosters, and we did not add HIV testing to the cost. We assumed that four booster doses after the primary first-year series were necessary to confer one decade of protection. For catch-up vaccination scenarios, booster eligibility was limited by the age range of the vaccinee. We did not, at this time, model the impact of more frequency booster doses.

### Nested hierarchy of vaccination

Motived by notion of a cascade of prevention [15], we conceptualized a nested hierarchy of vaccination using a mathematical model. We simulated a series of vaccination scenarios, sequentially incorporating different limitations of the vaccine and its roll-out. For each limitation, we quantified the decline in population-level impact, as measured by the percent reduction in new infections relative to a base case scenario without vaccine between 2027 and 2047 for a population aged 15-49. Since we did assume neither scale-up of ART coverage nor use of oral pre-exposure prophylaxis in the time period under investigation, percent reduction in new infections represents an upper bound relative to the maximum number of new infections prevented, i.e. more optimistic assumptions on ART scale-up would result in less infections prevented (data not shown).

First, we considered an idealized vaccine, providing complete protection (indefinite, 100% efficacy) without waning and available for use in 2027, even though we recognize the impracticality of such a scenario. In that scenario, we simulated a catch-up vaccination campaign among all men and women age 18-35 in 2017 at full coverage (100%), followed by vaccination of all 18-year-olds in subsequent years until 2047. Second, we simulated a more gradual vaccination, assuming a five-year linear scale-up starting in 2027 before reaching full coverage by 2032. Third, we considered a vaccine with limited duration of complete protection, assuming 10 years of full efficacy (100%), after which vaccine recipients were no longer protected. Next, we simulated a vaccine with partial efficacy that varied over 10-year protection period, averaging 50% over the first 2 years (including the first year of intensive vaccination) and falling to 15-30% over the next 8 years depending on booster frequency. Finally, we considered more realistic coverage levels of 60, 30, and 10% at various return rates for booster vaccination.

## 3 Results

Modeling results (see Figure 1) suggest that an ideal vaccine available in 2027 and covering all 18-35 year olds could prevent 89% of all infections occurring in the South African population aged 15-49 over the 2027-2047 interval. A more realistic 5-year ramp-up to reach full coverage entails a modest drop in impact to 79%, while limiting protection durability to 10 years results in a further drop to 66% infections averted. The second large reduction in intervention impact (from 66% to 20%) is due to assumptions of partial and waning efficacy. In total, factors related to vaccine durability, scale-up and efficacy result in a difference of 69% in vaccination effectiveness. Decreasing coverage from 100% to 60% will further attenuate the epidemic impact to just above 14% while problems with retention to the series of boosters will prevent as few as 11% of cumulative infections. In our most pessimistic scenario, assuming 10% coverage and 50% booster attrition, the epidemic impact drops down to only 3% of infections averted.

**Fig. 1.**
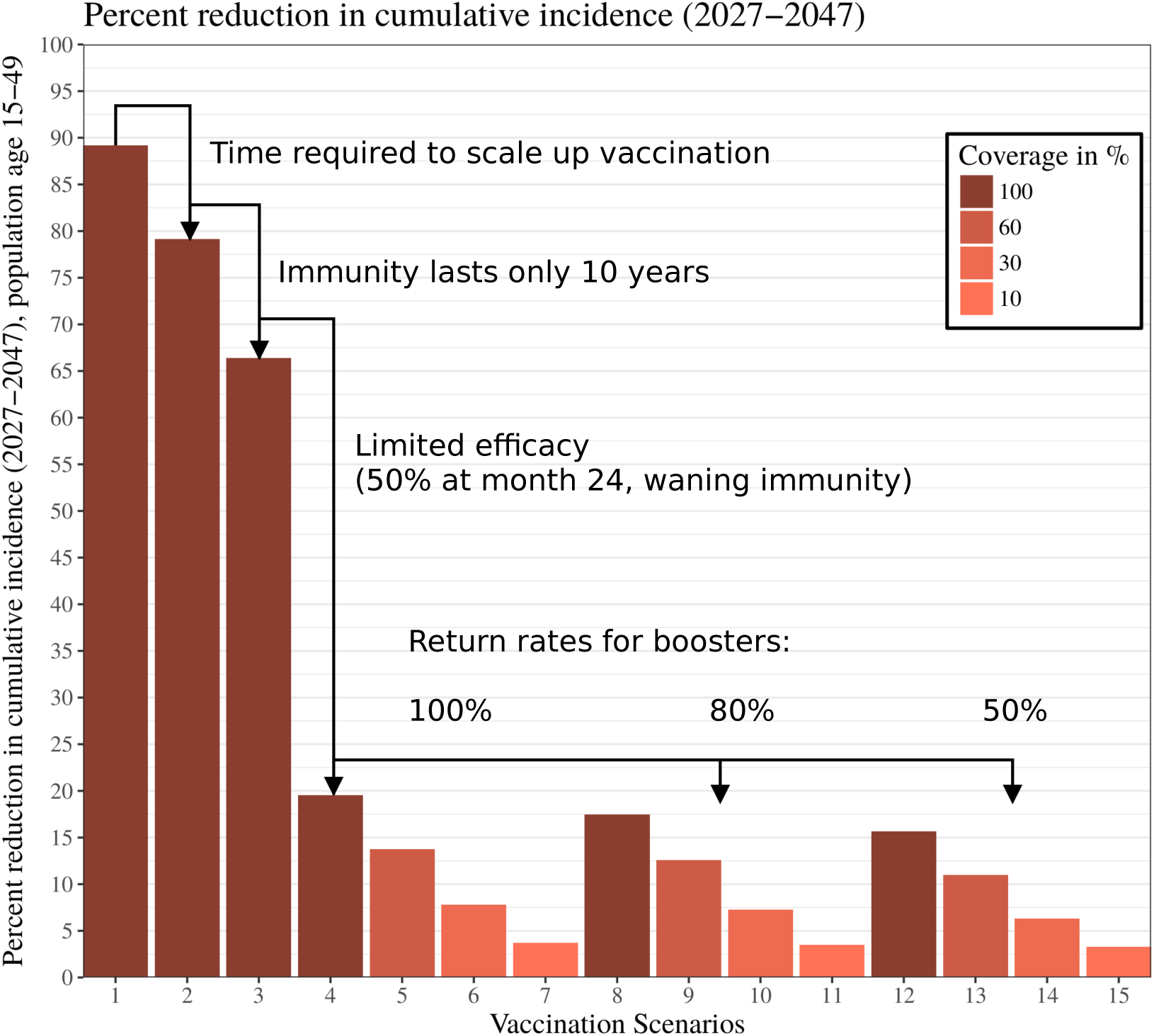
Percent reduction of new infections in a nested hierarchy of HIV vaccination: Vaccinating everybody between ages 18 and 34 with a perfect vaccine (sterile immunity, no waning) would prevent 89% of new infections within the core of the sexually active population (men and women aged 15-49). Under sequential limitations such as vaccine availability, limited durability or efficacy impact would decline to as low as 20%, despite full vaccination coverage. Decreasing return rates for boosters of the partially effective vaccine and limited coverage will further lower the impact to only 2% of infections prevented.

**Table 1.** Nested hierarchy of vaccination scenarios

| scenario | coverage | scale-up | efficacy | durability | waning | return rate<br>for booster |
| --- | --- | --- | --- | --- | --- | --- |
| 1 | 100% | 0 years | 100% | 20 years | no | NA |
| 2 | 100% | 5 years | 100% | 20 years | no | NA |
| 3 | 100% | 5 years | 100% | 10 years | no | NA |
| 4 | 100% | 5 years | 50% | 10 years | yes | 100% |
| 5 | 60% | 5 years | 50% | 10 years | yes | 100% |
| 6 | 30% | 5 years | 50% | 10 years | yes | 100% |
| 7 | 10% | 5 years | 50% | 10 years | yes | 100% |
| 8 | 100% | 5 years | 50% | 10 years | yes | 80% |
| 9 | 60% | 5 years | 50% | 10 years | yes | 80% |
| 10 | 30% | 5 years | 50% | 10 years | yes | 80% |
| 11 | 10% | 5 years | 50% | 10 years | yes | 80% |
| 12 | 100% | 5 years | 50% | 10 years | yes | 50% |
| 13 | 60% | 5 years | 50% | 10 years | yes | 50% |
| 14 | 30% | 5 years | 50% | 10 years | yes | 50% |
| 15 | 10% | 5 years | 50% | 10 years | yes | 50% |

## 4 Discussion

This modeling analysis of a nested hierarchy of vaccination clearly highlights three dominant reasons for the limited population-level impact of a partially effective HIV vaccine. Other important sources of limitations in the vaccination impact are the anticipated efficacy and durability of the pox-protein vaccine regimen as originally designed. We simulated 50% efficacy at month 24 as an illustration, but we hope that efficacy in HVTN 702 will be higher due to the modified regimen compared to RV144. Our analysis suggests that optimizing the efficacy of a broadly used vaccine should be a continuous process because of its critical contribution to the vaccination impact. Maximizing coverage has been rightly in the focus of all effectiveness analyses since it is likely to pose a challenge for a vaccine candidate with such a complex and lengthy dosing schedule. However, its ability to improve vaccination impact is already limited by the speed of vaccine roll-out and imperfect efficacy. Timely scale-up of manufacturing capacities, improved immunogenicity, and reassessing risk-benefit considerations for target populations with high risk profile during the licensure process could help to overcome the major obstacles to population-level impact identified in this analysis.

Model-based estimates of the impact of pox-protein-like vaccines are often met with disappointment. This analysis aims to clarify why the absolute impact of a partially effective multi-dose vaccine with limited durability is likely to be modest. Though worthwhile to develop and make available, such a vaccine is unlikely to reverse the course of the HIV epidemic. Nonetheless, it would provide a critical milestone for the development of a highly effective, durable, and scalable next-generation vaccine, which would have tremendous impact in combating the HIV epidemic.

## Acknowledgements

We acknowledge productive discussions with Andrew Phillips (University College London), Peggy Johnston, Silvija Staprans, Nina Russell and Geoffrey Garnett (all Bill & Melinda Gates Foundation).

CS, AB and PAW would like to thank Bill & Melinda Gates for their active support of this work and their sponsorship through the Global Good Fund.

DTD was supported through the Bill & Melinda Gates Foundation Award Number OPP1110049.

